# Singing and moving to the rhythm: song-entangled gestures in a vocal learning songbird

**DOI:** 10.64898/2026.08.13.744629

**Authors:** Emma Slupik, Ethan Ouyang, Elizabeth Joffrey, Spencer Kelly, Wan-chun Liu

## Abstract

When humans speak, we use rhythmic hand and head gestures that are closely coordinated with the temporal structure of speech to emphasize words and phrases. Similarly, human singing is often accompanied by rhythmic body movements that are aligned with the timing and prosodic structure of the vocal sequence. This rhythmic synchronization of bodily gesture with vocal production is thought to be a shared feature of vocal-learning species. How do these two sensorimotor systems develop, coordinate, and synchronize with precise timing to support multimodal communication? The mechanisms underlying this rhythmic entrainment remain poorly understood, and no established animal model to date captures the human combination of speech and co-speech gesture. Here, we show that a vocal-learning songbird, the zebra finch, has evolved song-entrained head gestures. These co-song gestures rhythmically align with acoustically complex song syllables and are developed and produced independently of other innate, stereotyped, song-entangled courtship displays. Even when the song remains largely the same, co-song gestures can be dynamically modulated in rhythm, form, and/or extent across different social contexts. The rhythmic alignment requires auditory feedback, is under the control of a premotor song nucleus, and gradually develops during the sensitive period of vocal learning. Females respond differently when songs and co-song gestures are misaligned, suggesting a social function. This dynamic modulation of co-song gestures provides a behavioral window into brain and cognitive states, making the zebra finch a promising model for understanding the mechanisms underlying rhythmic synchronization of multi-sensorimotor systems and the origin and evolution of co-speech gestures in humans.

## Introduction

Theories of language suggest that human speech evolved from–or co-evolved with–gestural communication systems in our evolutionary past [1, 2]. Co-speech gestures come in many different forms and rhythmically synchronize with specific aspects of speech to reveal cognitive, motivational, or emotional states of the human mind [3, 4]. Similarly, human music comes with a beat, and singing with rhythmically matched body movements helps the singer to regulate vocal rhythm internally and to highlight prosodic structure for the listener [5, 6]. A disruption to this rhythm synchronization impairs communication [7] and is evident in various developmental disorders [8–10]. How do these two sensorimotor systems develop, coordinate, and synchronize with precise timing to support multimodal communication and enhance social function? The neural mechanisms that underlie rhythmic entrainment between speech and co-speech gesture– or more broadly, between learned vocal and non-vocal motor control are poorly understood, and prior research has focused much more extensively on the rhythmic coupling between gestures and auditory perception of speech or music —in humans or in animals, such as parrots bobbing their heads to external musical beat. The sophisticated multi-sensorimotor integration and rhythmic synchronization involved in the planning, production, and perception of speech and gesture are likely served by a complex cortical-subcortical network [11–13].

While gestures have been well-documented in non-human species, there are currently no established animal models to explore learned vocal and non-vocal gesture synchronization and its underlying neural circuits. Songbirds are among a few groups of animals that have evolved rhythmic perception, production, and vocal learning [14–16]. As most previous birdsong research examined the rhythm of the song temporal structure [17, 18], auditory perception [16, 19], and the biomechanical production (i.e., respiratory or neural muscular control of the song) [20, 21], it is unclear whether there are non-vocal body gestures that rhythmically and precisely match temporal structure of the song through auditory feedback. Here, we show a vocal learning songbird, the zebra finch, have evolved song-entangled body gestures that are rhythmically synchronize with temporal structures of song.

## Results

### What are song-entangled gestures in zebra finches?

Co-song gestures in the zebra finch are rhythmic head movements that accompany song motifs. These gestures are primarily lateral head rotations dominated by yaw (rotation about the vertical axis, **Fig. 1A, Supplementary video #1**). Across individual birds, gestures occasionally consist of tilted movements, such as rotations dominated by roll (rotation about the longitudinal/forward axis) and pitch (rotation about the transverse axis). Importantly, like human co-speech gesture, the zebra finch’s co-song head gestures are consistently present across different social contexts, whether a male sings in isolation, in a colony, or during various social interactions. For example, a socially isolated male’s singing is accompanied by co-song head gestures while he stands still, without moving other body parts (**Supplementary Video #1**). These head gestures are incorporated into, but independently developed, produced, and controlled from the innate, female-directed, song-entangled courtship gestures of zebra finches (such as beak wipes, rhythmic hopping, pivot dancing, and fluffed-up head posture) [22–24].

**Figure 1.**
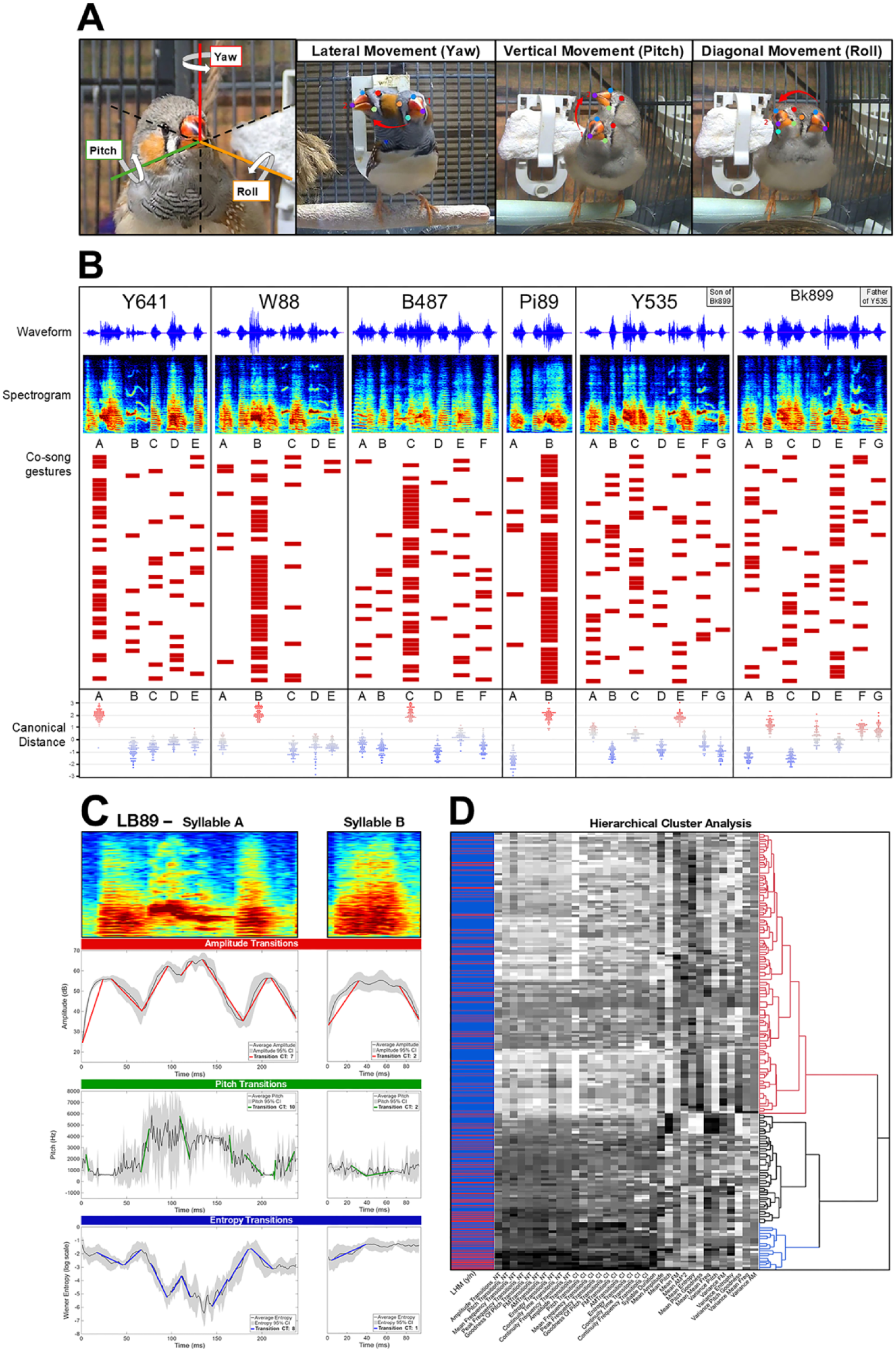
A) One of the most common co-song gestures: lateral head rotations dominated by yaw. Occasionally, adult males also display non-lateral head rotations while singing, such as movements that are predominantly pitch (vertical rotations) or roll (diagonal rotations). B) Idiosyncrasies in co-song gestures show alignment with more complex song syllables in each motif. Examples of co-song gestures from 6 male finches when singing alone (50 motifs per bird, alphabetical order represents the sequence of syllables for each song motif); each red square represents one lateral head rotation. Each row represents one motif. Lower panel. The most gestured syllables per motif in each bird have significantly more distinct acoustic features (red dots) compared to syllables with fewer gestures (blue dots). Canonical distance measures the multivariate acoustic distance among syllables per motif in each bird. C) Accumulative acoustic transition counts of syllables predict co-song gesture affiliation. Each colored line segment represents a significant transition in an acoustic feature during the time interval of a syllable. Transitions defined as significant have endpoints with non-overlapping 95% confidence intervals (CIs, highlighted gray). LB89- syllable A (left) has 60 more significant transitions than LB89-B (amongst all 10 tested acoustic features) and is gestured on more frequently (80% gesture rate). D) Dendrogram of hierarchical cluster analysis using 34 acoustic feature descriptive statistics (transition counts, mean, and variance values) from 9026 syllables to categorize complex syllables (darker blocks represent syllables with higher values of an acoustic feature) that are associated with more song gestures (red lines represent syllables that were gestured on, blue represent syllables that were not gestured on).

Co-song gestures do not randomly occur on song syllables (Discriminant function analysis, lateral head movement quantified from 9026 syllables of 30 adult males, Λ = 0.907, *X^2^* = 751.08, *p <* 0.001). Instead, co-song gestures are aligned with a specific song syllable within each motif (temporal matching in tens of milliseconds, **Fig. 1B**). These syllables are characterized by longer syllable duration and higher variance of acoustic features. Depending on individual birds, some of the strongest predictors for co-song gestures are the syllables with higher variance of entropy, pitch, and amplitude modulation (Discriminant function analysis, *F* = 13.800-233.753, *p <* 0.001), suggesting that co-song gestures are more likely to be aligned with more acoustically complex syllables within each motif.

### Rhythmic alignment with complex syllables

To quantify gesture-associated syllable complexity, we calculated the number of transitions– increasing or decreasing intervals between local extrema–of acoustic features within each syllable (**Fig. 1C**, **Supplementary Figure 1**), as a higher number of acoustic transitions within each syllable presumably requires more elaborate and complex neuromuscular coordination [17]. We found that adult males produce significantly more lateral head gestures on syllables with more acoustic transitions. This relationship exists between both total transition accumulation (Pearson’s Correlation, *r* = 0.716, *p <* 0.001) and when filtering for significant transitions that are robust to rendition-to-rendition acoustic variation (Pearson’s Correlation, *r* = 0.618, *p <* 0.001, **Supplementary Figure 1**). Across all birds, while all acoustic transition counts were significant predictors of gestures (Discriminant function analysis, *X^2^* = 874.617; *p <* 0.001), some of the strongest predictors for co-song gestures were the number of total transitions in continuity frequency, pitch, goodness of pitch, and the number of significant transitions in amplitude and entropy per syllable (Discriminant function analysis, *F* = 517.053-762.953; *p <* 0.001, *n* = 30 birds). As zebra finches are able to discriminate extremely small changes in the fine acoustic structure of syllables, there is the potential for information such as sex, motivational/emotional state, and identity to be transmitted via transitions in acoustic features of these more complex syllables, which may be emphasized via co-song gesture alignment [25, 26].

Although syllable duration is one of the strongest predictors of co-song gesture (*F* = 734.36, *p <* 0.001), we rule out that co-song gesture occurrence is solely associated with more acoustically complex syllables due to the increased chance of landing on longer syllables. After controlling for the syllable duration for each syllable, the higher ratio of syllable feature transitions/syllable duration remains significant in predicting gestured syllables (*X^2^* = 286.28, *p <* 0.01). Furthermore, our deafening experiment shows that co-song gestures fail to align with more complex song syllables after deprivation of auditory feedback (see below **Fig. 4**).

Co-song gestures are highly idiosyncratic across birds; their degree of alignment with syllables varies by individual. Therefore, each bird has a unique gesture rhythm (**Fig. 1B**). For example, even when a male tutee precisely imitates the song of its father tutor, the rhythmic patterns of co-song gesture of the tutee can differ from those of its father tutor, even if the tutor song was precisely imitated (Y535 vs. Bk899 in **Fig. 1B**, in each of these two birds, there are multiple syllables per song motif that show higher score of syllable complexity).

### Co-song gestures are social context dependent

In humans, co-speech gestures are significantly influenced by social contexts [27]. When singing alone (undirected song) or under social isolation, we found that male zebra finches make robust co-song head gestures. However, in the presence of a female, males produce more frequent lateral (yaw-centric) co-song gestures, especially for gestures aligned with the complex syllables (Mann-Whitney U test, *U* = 16, *p* = 0.003, *n* = 11 birds, **Fig. 2B**). While these head gestures are often accompanied with well-documented innate courtship displays in female directed contexts– including beak wipes, stretching and puffed-up body posture, hopping and moving around–co- song gestures are present in any context, are independently developed, and may be separately controlled from courtship displays (see below). In some males, the rhythms of gesture across song motifs also change or intensify in female-directed contexts (see examples in **Fig. 2A**), perhaps due to the increased song tempo and to better align with the body movement of courtship gestures.

**Figure 2.**
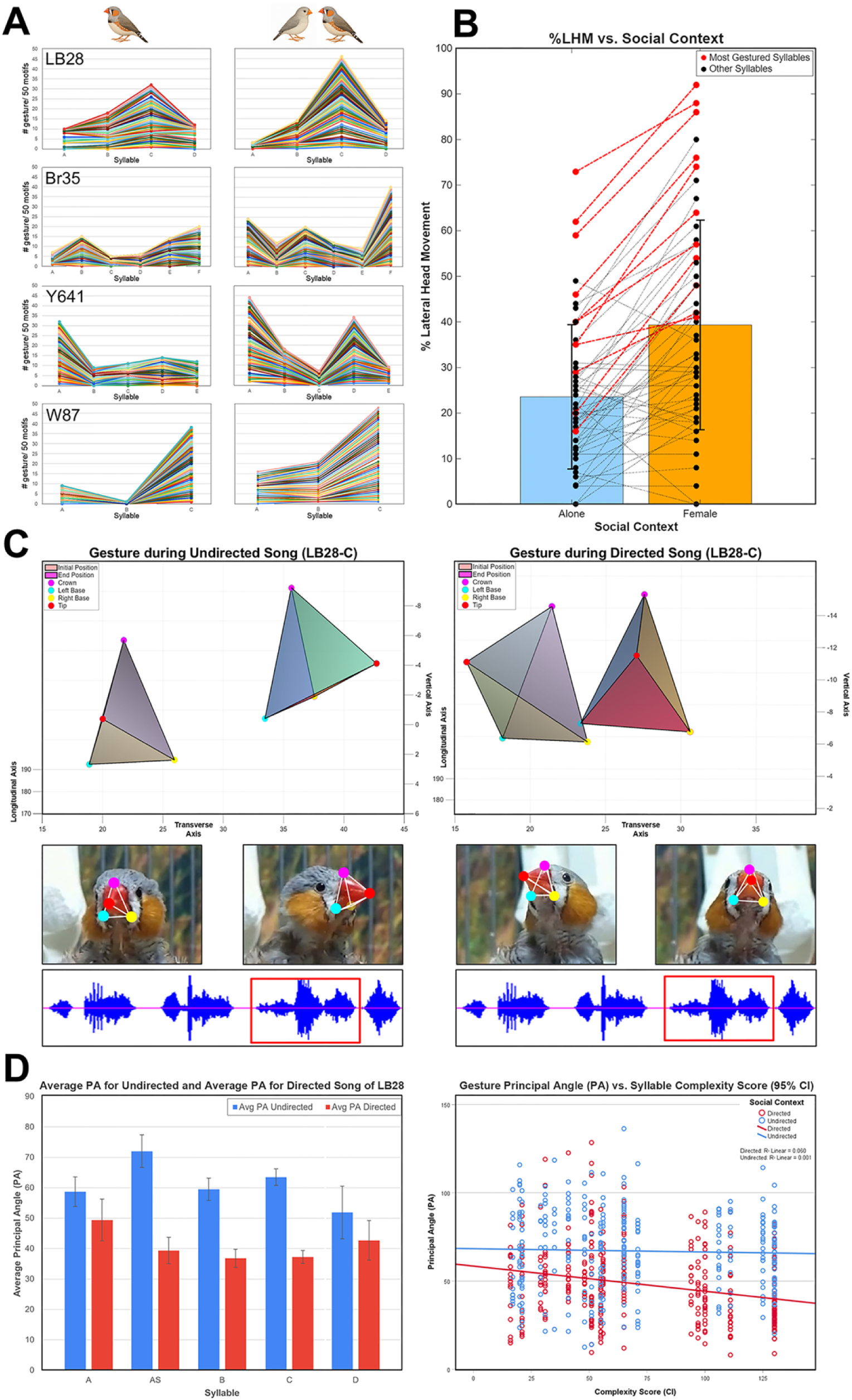
Rhythmic movement, frequency, and intensity of co-song gestures are social context dependent. **A)** Four examples of rhythmic changes of co-song gestures when a male sings undirected song (left panel) vs. song directed toward a female (right panel). Each colored line represents one motif. **B)** When singing to a female, the frequency of co-song gestures increases (%), especially during the most complex syllables (red dots and lines) of each motif per bird (*n* =11 birds). **C)** When performing undirected songs, birds gesture with a greater degree (PA) compared to when performing female-directed songs. **Left panels.** Example of the 3D reconstructed beak positions for the initial and end positions of a lateral co-song gesture during undirected song (LB28-C syllable). Using an implementation of the Kabsch Algorithm, the PA for this movement was 69.98° (contributions: 99.9% yaw, 3.8% pitch, 1.0% roll). **Right Panels.** Example of 3D reconstructed beak positions for a co-song gesture during female directed song (LB28-C). PA = 37.77° (contributions: 98.3% yaw, 9.8% pitch, 16.4% roll). **D) Left Panel** A syllable-by-syllable comparison for LB28 of the average gesture PA during undirected (blue) and directed (red) songs. **Right Panel.** As syllable complexity increases (x-axis), the PA of gestures on the syllable (y-axis) decreases more drastically between undirected (blue) and directed songs (red).

Additionally, results from our 3D model further reveal the magnitude of gesture movement changes under different social contexts. Gesture magnitude is measured via principal angle (PA); PA quantifies the angular magnitude of rotation required to map the initial head pose to the end head pose. During undirected singing, the magnitude of gesture movement is significantly larger than directed singing (undirected PA, *M* ± *SD* = 67.00 ± 22.58; directed PA, *M* ± *SD* = 48.32 ± 21.34, linear mixed model, main effect of social context on PA, F(1, 640.12) = 6.94, p = 0.009, **Fig. 2C**). Furthermore, there is a significant interaction between social context and syllable complexity on gesture PA (linear mixed model, *F*(1, 649.14) = 9.35, *p* = 0.002); as males sing directed song, the magnitude of gesture movement (PA) significantly decreases while singing the more complex syllables, a decrease not observed in undirected song (**Fig. 2D**). These results show that the intensity and rhythm of a co-song gesture can be dynamically modified under different social contexts, even when vocal acoustics remain largely the same. This suggests that beyond a simple biomechanical function, co-song gestures may be driven by emotion, motivation, or social goals.

### Gradual development of co-song gestures

In human language acquisition, infants make uncoordinated gestures before and during the early onset of babbling, and then gradually align them with acquired speech [28–30]. We tracked the development of co-song gestures over the sensitive period of song learning in juvenile zebra finches (*n* = 5 males) and found a significant interaction effect between gesture/rotational type and developmental period (linear mixed model analysis, *F*(21, 77.62) = 2.59, *p* = 0.001).

Juveniles in the early sensorimotor learning phase, or subsong stage (39-40 days post-hatching, or dph), display more non-lateral head movements; roll rotations are significantly elevated from subsong and adult gestures during early plastic song (49-50 dph) (post-hoc, Bonferroni between early plastic and subsong, *p =* 0.047, **Fig. 3A, C**). Gestures then gradually lateralize and resemble a stereotyped adult gesture pattern when the sequence of song syllables (or motif) stabilizes toward the end of the sensitive period (63-64 dph). Lateralization significantly increases up until young adulthood (approximately 120 dph, Bonferroni adjusted, *p* < 0.001, **Fig. 3A**) before stabilizing in older adulthood (1 year of age or older, **Fig. 3C**). Similarly, the number of gestures produced per syllable remains high during the sensitive period of song learning but slowly reduces at 65 dph and then significantly decreases at 120 dph or older (ANOVA, *F =* 3.72, *p <* 0.01, **Fig. 3B**). While gradual lateralization is observed, its degree and timescale are idiosyncratic, as movement rates largely differ by bird across each time point (mean CV across rotational type = 52.3-88.5%). This individual variability among birds might be associated with individual specific song learning strategies [31]. These results suggest that the stability of rhythmic matching of co-song gesture with complex syllables may gradually develop after the emergence of song syntax (stereotyped sequence of song syllables), while the development of the innate, song-entangled courtship gestures (hopping, beak wiping, head/body fluffing postures) occurs as early as 30-35 dph in juvenile finches [32].

**Figure 3.**
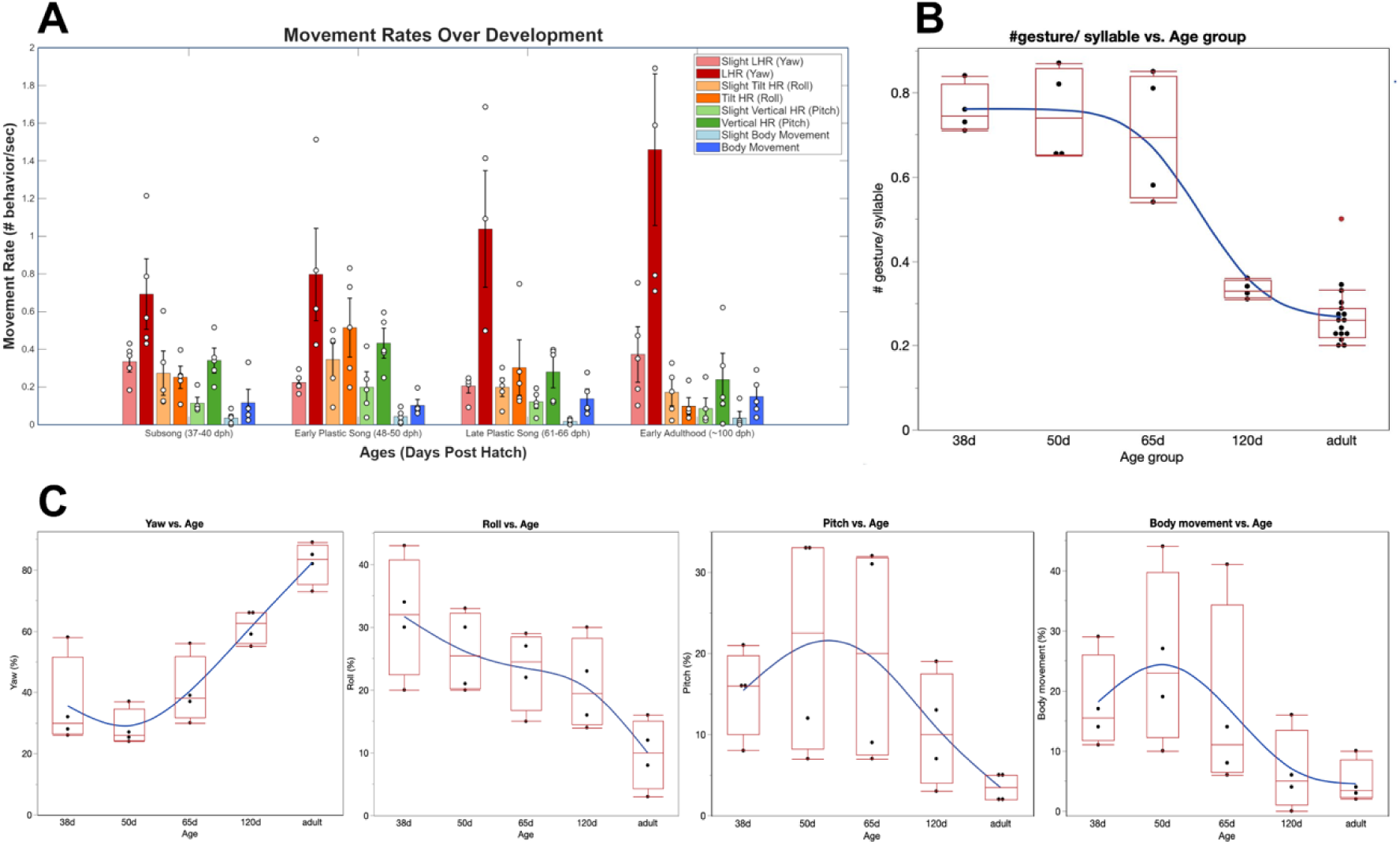
Co-song gestures become more stereotyped throughout song development. **A)** Juveniles in early sensorimotor development display more non-lateral head rotations such as vertical, pitch-dominated rotations and diagonal, roll-dominated rotations. Head and body movement rates were calculated for various timepoints in song development (*n* = 5 males). Juveniles in early development also produced more slight head movements (head rotations less than 45°). **B)** During early development, juveniles produced a higher number of co-song gestures when singing. The gesture frequency decreased as adults. **C)** Over the course of development, the percentage of yaw rotations (lateral head movement) gradually increased toward adulthood, while pitch (vertical head movements), roll (diagonal head movements), and body movements gradually decreased.

### The alignment of song gestures and complex syllables requires auditory feedback

The gradual stereotyping of co-song gestures during song development suggests that the rhythmic alignment of co-song gestures with complex syllables may require auditory feedback, a necessary trait for vocal learning species, and one suggested in human study of co-speech gestures [33]. We tested this hypothesis by conducting bilateral deafening of adult males (*n* = 5 males, > 1 year old). Approximately two months post-deafening, the song remained mostly intact, and yet the co-song gestures no longer aligned with complex syllables; gestures were more randomly distributed across all syllables and therefore had no clear rhythm (*X^2^* =8.34, *p*=0.304, **Fig. 4 A, B)**. Interestingly, post-deafened birds also produced more “roll” and “pitch” head gestures and more “slight” lateral movement, reminiscent of juvenile developing gestures (**Fig. 3**). Rhythmic matching of song and co-song gesture therefore requires the integration of auditory input (hearing the bird’s own song) and motor output (co-song gesture). Much like the maintenance of learned songs, auditory feedback may provide a mechanism for error correction of misaligned songs and gestures.

**Figure 4.**
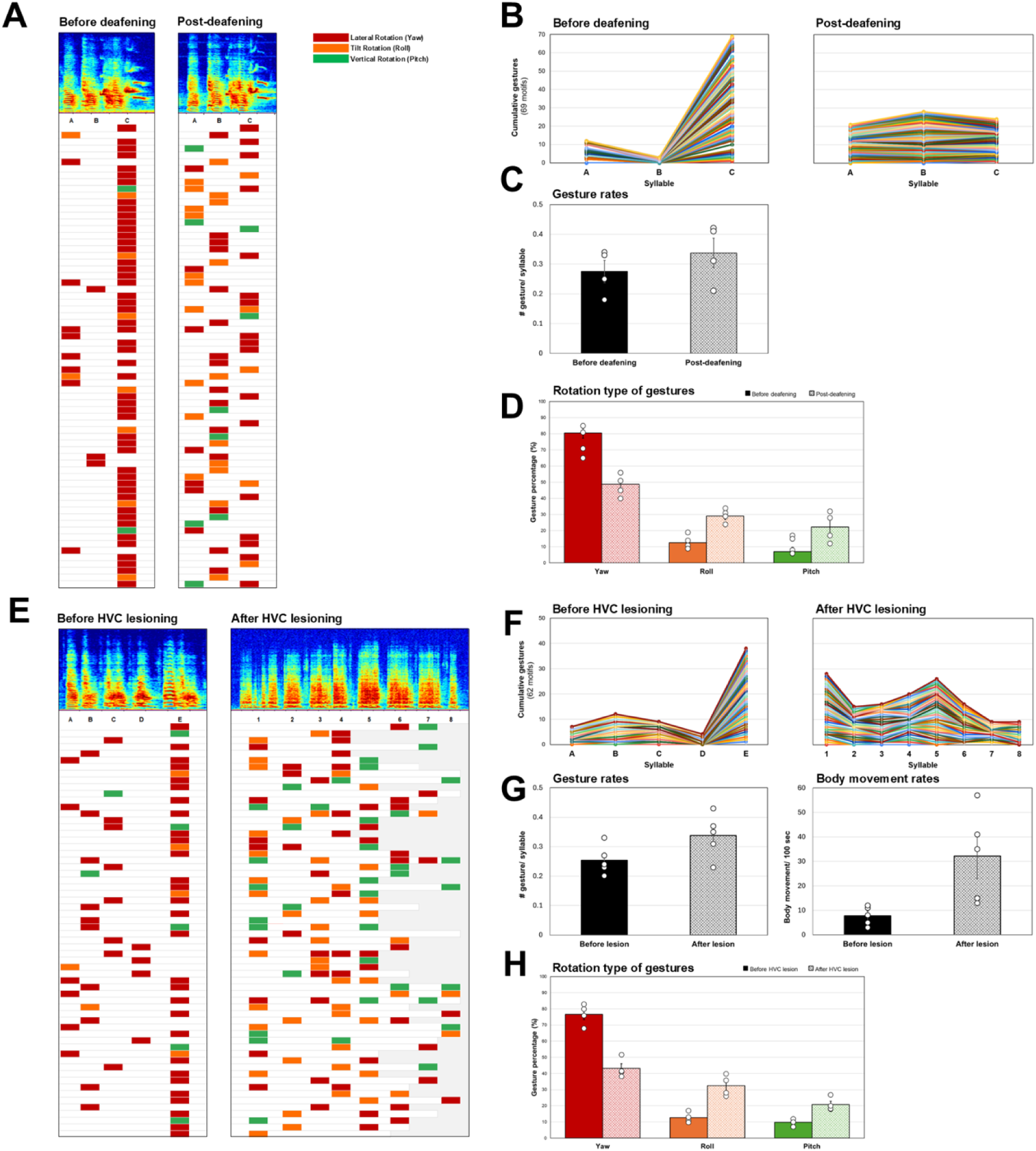
Bilateral deafening and HVC lesioning disrupts the rhythmic alignment of co-song gestures with complex syllables. **A-B)** An example to show that co-song gestures of an adult male, W87, were aligned with a complex syllable “C” before deafening. Two months after deafening, his song remained mostly intact, but his co-song gestures were more randomly distributed across all syllables, suggesting a dependence on auditory feedback. **B)** The changes of rhythmic matching of song gestures after deafening of W87. **C-D)** Post-deafened birds had more roll and pitch gestures and produced more gestures overall (Mann-Whitney U test, *U* = 2.48, *p <* 0.05, *n* = 5 birds). **E-H)** HVC lesioning disrupts the rhythm and production of co-song gestures**. E)** An example to show that co-song gestures of an adult male, B487, were aligned with a complex syllable “E” before HVC lesioning. After lesioning (1-2 weeks post-lesioning), the song syllables were gone, and co-song gestures were randomly distributed across syllables in each song bout. **F)** The rhythmic changes of HVC-lesioned co-song gestures. **G)** Post-lesioned birds have significantly more head gestures and body movements than before lesioning and **H**) produced more roll and pitch gestures (*n* = 5 birds). Type or paste table title here. Paste table below the title.

### The rhythm alignment of co-song gestures involves a premotor song nucleus

The ability to rhythmically produce gestures in synchrony with perceived speech and music relies on a cortical circuit [16, 19, 34]. We hypothesize that the entrainment between song production and non-vocal gesture movement also requires cortical control. To test this hypothesis, we conducted bilateral lesioning of a cortical song nucleus, the HVC, and examined its effect on the rhythmic matching between song and co-song gestures (*n* = 5 birds). HVC is known to be a generator of song rhythm[18]. Does HVC involve in the rhythm alignment with co-song head movement? Bilateral HVC lesioning immediately disrupted the rhythm and production of co-song gestures (**Fig. 4 E-H**). In post-lesioned birds, the acoustic structure of learned syllables and syntax disappeared (resulting in subsong-like song [35]), and co-song gestures were randomly distributed across syllables in each singing bout. Additionally, post-lesioned birds produced more “roll” and “pitch” gestures and had significantly more body movements than before lesioning (*n* = 5 birds; Mann-Whitney U test, *Z* = -2.61, *P* = 0.009, **Fig. 4G**), reminiscent of the random gesture movements produced during song development (**Fig. 3**). These results show that HVC lesioning not only abolished song production but also disrupted the rhythmic synchronization between song and head gesture movement, which suggests that the rhythmic entrainment of song and co-song gesture requires the involvement and coordination of cortical song circuits.

Post-HVC-lesioned birds produced atypical, arrhythmic co-song gestures. This is in striking contrast to other female-oriented co-song courtship displays in zebra finches which are under the control by a midbrain nucleus [24], and persist even after HVC lesions [24]; and they do not require auditory feedback [36].

### Social communicative function of co-song gestures

The robust, non-random presence of co-song gestures and their sensitivity to social context suggest that co-song gestures have a social communicative function, perhaps to enhance, modulate, or amplify vocal signals (song) toward their receivers under various social contexts. We thus hypothesize that conspecific signal receivers would be able to recognize the alignment or misalignment between syllables and gestures. To test this hypothesis, we conducted video playback experiments by simultaneously playing back audio-delayed videos (1 sec of audio-video mismatched) and normal, audio-video matched videos (both from the same recordings of the same bird when singing alone, *n* = 7 males recorded), toward adult females. We found that adult females (*n* = 8 virgin females) spent significantly more time on the perch near the normal, synchronous playback video (**Supplementary Figure 2**, *n* = 8 females, Mann-Whitney U test, *U* = 20, *Z* = -3.58, *p <* 0.01), suggesting the females can differentiate between matched and mismatched co-song gestures, similar to humans’ sensitivity to the temporal alignment of speech and beat (rhythmic) gestures (*9, 14*).

## Discussion

Co-song gestures that we observed in male zebra finches are produced, developed, and controlled independently of female-directed, innate, stereotyped song-entangled displays [22–24]. In contrast to these female-oriented courtship displays, co-song head gestures are more like human co-speech gestures, as they are produced and are dynamically modulated in various social contexts. Like humans, co-song gestures gradually develop and align with complex song syllables over the course of song learning; and their rhythmic matching relies on the engagement of auditory feedback (**Fig. 4**). Moreover, lesioning the HVC not only abolishes learned song but also severely disrupts the rhythm and forms of co-song head gestures– yet innate courtship displays persist after HVC lesioning (20) and after the deprivation of auditory feedback [36]. The context-dependent nature and dynamic modulations of co-song gestures mirror context-dependent neural variability in the basal ganglia network [37], potentially offering a more sensitive behavioral indicator of underlying brain and cognitive states.

The involvement of a premotor song nucleus and auditory feedback for the rhythmic synchronization between song and co-song gestures in zebra finches suggests a close circuit connection between learned vocal and non-vocal motor gestures. In songbirds, the learned-song circuitry is remarkably vocal-specialized. This specialized “song system” is dedicated to vocal learning, vocal production, and the biomechanical production and rhythms of song structure. For example, HVC is known to be a main generator of the song structure [18], and HVC neuron bursting gradually shifts and slows down its rhythms over the course of song development [38]. Similarly, here we show that co-song gestures are produced more frequently and randomly during early song development, and gradually slow down and become stereotyped after the emergence of song syntax. Bilateral HVC lesions not only demolish song structure and its rhythm, but the temporal coordination between co-song gestures and syllables is also lost, resulting in significant disruptions to the rhythm and forms of head movement (**Fig. 4**). Moreover, in post-deafened birds, gestures lose their alignment with specific syllables and become arrhythmic, even though those syllables persist, suggesting syllable-gesture coupling requires feedback-dependent maintenance. Together, these results suggest that production of song-locked gesture rhythm relies on the auditory feedback of the bird’s own song and is coordinated with HVC. There may exist an integration of overlapping or adjacent motor/auditory circuits that favors the evolution of rhythm synchronization between speech production, perception, and aligned non-vocal body gestures [13, 39, 40].

The rhythmic matching between complex syllables and co-song head gestures is reminiscent of how co-speech beat gestures in humans emphasize particular words or phrases within larger spans of discourse [41] –even altering the acoustic prominence of speech [42–44]. Moreover, the dynamic modulation of co-song gestures in zebra finches, suggested from our 3D analysis (**Fig. 3 C-D**), may help receivers to better perceive and identify a signaler’s emotional or motivational state. For example, idiosyncratic gestures with larger magnitude and less frequent lateral movements are commonly observed in an “undirected” context, which may enhance individual recognition from small social gatherings during non-breeding season [45, 46]; while smaller, intense, and more stereotyped lateral gestures, emphasizing complex syllables, are used in female-oriented contexts, perhaps allowing males to fast flash and “highlight” their orange cheek patches in a close range to impress females, which loosely resembles a deictic “pointing” gesture. Although non-human animals commonly use visual displays to draw attention to particular body parts, very little is known about whether such actions are temporally coordinated with specific elements of learned vocal signals. The study of context-dependent co-song gestures in zebra finches or other vocal learning songbirds may shed light on the origin and evolution of complex co-speech gestures in humans [1, 47].

Most prior research on co-song gestures in songbirds examined the female-directed courtship display [48, 49]; and the body gestures accompanied with song are thought to be rooted in biomechanical adaptation for respiration to enhance signal transmission [50] (for example, the head/body stretching or bowing of co-song display in brown-headed cowbirds [51, 52]). Male zebra finches learn and produce only one single song type. And yet, their learned song serves multiple social functions beyond the courtship display to attract females [46]. Even when the vocal aspects of the song remain largely the same, each adult male can dynamically modulate co-song lateral head gestures in different rhythms, magnitudes, frequencies, and under different social contexts; moreover, these lateral head movements appear to have no clear biomechanical connection with the respiration or production of song. In this way, adult males may use co-song gestures to deliberately emphasize complex song syllables, while also generating idiosyncratic rhythmic movement during the integrative production of song and co-song gesture (**Fig. 1**). Our results from the playback experiments show that the female can perceive desynchrony (mismatch in timing) between song and co-song gestures (**Supplementary Fig. 2**), further suggesting a social function of the co-song gesture. The idiosyncratic and rhythmic co-song gestures may allow individual finches who live in a complex social environment to efficiently communicate with one another by revealing their social and cognitive status or by providing an honest signal.

## Materials and Methods

### Animals

A total of 75 male and female sexually mature zebra finches (*Taeniopygia guttata*) from Colgate University’s animal facility were used for this study. All birds were kept on a 12:12 light cycle from 0800 to 2000, and had unlimited access to seeds, water, vegetables, egg supplement, and grit ad libitum. Thirty adult male zebra finches were used for the recording and behavioral coding analysis of co-song gestures. Each male selected was at least one year old, having reached sexuality maturity and song crystallization (occurring around 90 dph). Total song complexity and duration varied by bird, as did their breeding status and relation to other recorded birds; tutors were selected to examine whether idiosyncrasy in co-song gesturing patterns exists even with similar song structures. All treatments and experimental procedures were approved by the Institutional Animal Care and Use Committee’s (IACUC) at Colgate University (2425-13).

### Data collection

To record and quantify co-song gestures during singing, the undirected song of 30 adult male zebra finches was recorded between 2020 and 2025. In brief, adult male finches were individually kept in a semi-social environment, wherein they were kept in complete visual isolation and partial audio isolation from other birds. They were allowed to acclimate in chambers for at least 3 days before the start of the recordings. Cages were 25.5 × 30.5 × 33 cm in size, with a square plexiglass window (22 x 22 cm) on the front wall. Birds were kept under a L12:D12 day: night cycle (9 AM-9 PM). Video footage was obtained using WiFi-connected Reolink E1 Zoom or Reolink 4MP cameras placed at the front of the glass panel on the cage, aligned with a perch (**Supplementary Fig. 3**). For audio recording, condenser microphones (Audio Technica, AT801) were placed near the center of each cage and continuous audio recording occurred using Sound Analysis Pro (SAP 2011). A minimum of 50 song motifs were collected per bird, mostly in the morning hours.

### Behavioral coding of co-song gestures

First, we identified each bird’s song motif and syllables using Raven Lite: Interactive Sound Analysis Software (Version 2.0.5, Cornell, Ithaca) and examined the spectrogram and waveforms of multiple audio files of songs. Once a motif was identified, each song syllable of a motif was then categorized in alphabetical order. Individual syllables were identified if there was a minimum silent interval of 5 ms in the motif [53]. Within our samples, motifs ranged in syllable quantity from two (that is, syllables A-B) to nine (syllables A-I). Motif quantifications were verified with 1-2 additional coders before proceeding with behavioral analysis. Subsequently, the co-song gestures present in at least 50 motifs of each bird’s song (Mean ± SD = 57.67 ± 19.40 motifs coded) were analyzed using Behavioral Observation Research Interactive Software (BORIS).

Spectrograms were used to visually identify song syllables and motifs. The frame-by-frame click through feature, adjustable playback speed to slow down the video clips, and simultaneous visual of the waveform or spectrogram allowed for precise coding of head movements and the determination of which syllable the head movement occurred on. Separate behaviors were created to record head gestures occurring on different motif syllables, denoted by letter keys (e.g. “a” for a gesture on the A syllable). Once recorded, behavior events were double-checked to ensure each gesture occurred precisely on its recorded syllable (occasional skipping of the waveform visual was corrected for). Overall, 9,026 total syllables from 30 birds were recorded.

Behavioral results were reformatted to facilitate further analysis, and the gesturing frequencies per syllable were calculated, both using a custom MATLAB program (version R2025A, MathWorks). For coding analysis, two or three students and WL independently coded for the same bird for at least 10 motifs/ per bird, and the coding data was recorded if independent coders reached similar agreement.

### Song syllable analysis and statistical analysis

The acoustic features of song syllables were analyzed using SAP2011. For each of the 50 song motifs per bird, each syllable was selected, and the mean and variance values for eight acoustic features (amplitude, pitch, FM, AM^2^, entropy, pitch goodness, mean frequency, syllable duration) were collected using SAP. Fifty records of the same syllable were averaged to reduce the influence of natural variability in song, likely reflecting aspects such as motivational state, breeding status, hormone levels, temperature, or time of day. For statistical analysis, a multivariate, discriminant function analysis was conducted to determine whether co-song gestures rhythmically aligned with specific song syllables within each motif as opposed to occurring randomly throughout the syllables in each motif, and what combination of acoustic features best predicted whether a co-song gesture occurred on a particular syllable in the motif (SPSS Statistics version 29.0.2.0). SAS JMP (version 18.2.1) further allowed us to visualize the relative canonical distance among syllables in each motif, as an indicator of acoustic differences among syllables. This software also allowed us to establish and visualize a dendrogram from a hierarchical cluster analysis to identify the association between acoustic features of syllables and syllable-specific gestures. This same statistical test was later run with the inclusion of transition values for each sound feature once syllable complexity analysis quantified these transition counts (for both tests, *n* = 9026 syllables sampled from 30 adult male zebra finches).

### Complex syllable analysis

Using SAP, sound files for 155 distinct syllables (from the sample of adult male zebra finches previously recorded, *n* = 30) were analyzed for the following ten variables at each timepoint within a syllable interval: amplitude, pitch, mean frequency, peak frequency, goodness of pitch, FM, AM, entropy, continuity time, and continuity frequency. A minimum of five renditions of each syllable were sampled (787 total samples) to reduce the influence of natural variability in song and generate 95% confidence intervals (CIs) for each timepoint. These data were fed into a MATLAB model that generated an average interval plot for each syllable feature by resampling each sample so each is matched in duration (using the MATLAB “resample” function, samples were interpolated or decimated to match the average syllable duration; a FIR antialiasing lowpass filter was automatically applied by the function to prevent distortion during resampling). Local maxima and minima for each acoustic feature’s average line were calculated and intervals between them were considered as transitions. To adopt and compare multiple strategies, one program variation totaled all transitions at this point (termed: “No Threshold Transitions”), whereas the other identified significant transitions between local maxima/minima by selecting the transition closest in time between two points without overlapping CIs (a greedy algorithm approach was used).

Overlapping significant transitions were filtered out to prevent double counting; this strategy is referred to as “95% CI Transitions”; **Supplementary Fig. 1**). Both methods for counting acoustic transitions were positively correlated with lateral head gesture frequency (No threshold: Pearson’s Correlation, *r* =.716 *P<*0.001, correlation between significant transition counts and gesture frequency on a syllable; 95% CI: Pearson’s Correlation, *r* =.618 *P<*0.001). We performed a multivariate, discriminant function analysis in SPSS to determine whether transition counts of either method predicted the timing of co-song gestures.

### Development of co-song gestures

Five juvenile male zebra finches were recorded for the development study. Four breeding pairs were selected from the Colgate University bird colony and birds hatched in approximately mid-August 2025. Once the juveniles were close to independence (approximately 30 dph), juveniles and their song tutors were removed from the bird colony and placed in the recording chambers in pairs.

### Song and gesture recording

The undirected songs of 5 juvenile male zebra finches were recorded (audio and video) at regular intervals from 30 day post-hatch (dph) to early adulthood (∼120 dph), using condenser microphones (Audio Technica, AT801) and Reolink E1 zoom cameras. See the previous section for the recording environment (cage and recording setup; **Supplementary Fig. 3**). Timepoints during the sensitive period associated with rapid changes in song development were recorded on alternate days; the transition between subsong and plastic song (between approximately 40 and 50 dph) was closely monitored. Late subsong (38-40 dph), early plastic song (48-50 dph), and late plastic song (once song became largely stereotyped at 63-65 dph and onwards) were recorded every other day. As song learning has been shown to begin around 25 dph and require approximately 10 days of contact with a song tutor to create an accurate imitation of their song, adult tutors remained in the recording cage with the juveniles until 40 dph (*36*). After this time, juveniles were kept in visual isolation and partial audio isolation from other birds until song crystallization.

### Co-song gesture analysis

Using a revised behavioral coding scheme on BORIS, a variety of movements were quantified at four distinct developmental periods per bird: subsong (38-40 dph), early plastic song (48-50 dph), late plastic song (63-66 dph), and early adult song (∼120 dph). Each interval consisted of recordings from at least two separate recording days to reduce variability in movement due to exact time of recording or the bird’s motivational state, as birds were observed to be more reserved in movement in certain recordings and more excited in others, regardless of developmental stage. Movements identified during song included head pose rotation parameters and body movements; lateral head movements (yaw-dominated), vertical head movements (pitch-dominated), diagonal head movements (roll-dominated), and full body movements were recorded. “Slight”, visually distinct small movements, varieties of each movement type were also recorded. Thus, the movement behaviors recorded on BORIS included (each signified with a unique letter key): “Lateral Head Movement (Yaw),” “Slight Lateral Head Movement (Yaw),” “Diagonal Head Movement (Roll),” “Slight Diagonal Head Movement (Roll),” “Vertical Head Movement (Pitch),” “Slight Vertical Head Movement (Pitch),” “Full Body Movement,” and “Slight Full Body Movement.” To explore whether movement rates of different rotational types changed significantly over development, we conducted a linear mixed model analysis using SPSS

Statistics (version 29.0.2.0). To ensure normality of residuals and homogeneity of variance, movement rates were transformed using a square-root transformation. Using a first-order autoregressive correlation structure to analyze longitudinal data, AR(1), the model included movement behavior type and developmental period as fixed effects and random intercepts for birds and slopes for each time period, grouped by bird (a variance component (VC) covariance type was used). We used this random effect structure to account for different baseline movement rates of birds and individual differences in the timing of their development.

### Stereo vision recording of gesture analysis

Male zebra finches were recorded in a cage with plexiglass on the front side of the cage (**Supplementary Figure 4**). Video was then captured using two 12.3MP 477P Raspberry Pi Arducam cameras and Camarray HAT for stereo vision video capture. The cameras were placed edge to edge with a baseline distance of 38.0 mm. The cameras were then mounted to a camera mount that was attached to a metal base plate drilled into a table. For audio recording, a condenser microphone (Audio Technica, AT801) was placed near the center of the cage and was connected to an audio interface (Behringer U Phoria UMC202HD USB Audio Interface). The cameras and the audio interface were connected to a Raspberry Pi 4 (RPi4) to capture simultaneous audio and video.

### 3D reconstruction model for gesture analysis

The Arducam Cameras were calibrated using MATLAB’s stereo camera calibrator and a 10 x 12 checkerboard with a checker size of 8 mm. Videos captured from the stereo vision recording setup were split into two videos (left and right) and rescaled using FFMPEG. Each video was fed into a trained DeepLabCut mode [54, 55] to generate 2D coordinates for key points of the beak (crown, left base corner, right base corner, and tip) for both the left and right camera. These coordinates were then fed into a MATLAB model that treated the beak as a rigid body tetrahedron. The model performed triangulation in order to reconstruct the beak in a 3D space where the x-axis is the transverse axis, z-axis longitudinal axis, and y-axis is the vertical axis.

Each initial reconstruction was compared to a reference tetrahedron beak using an implemented Kabsch Algorithm. The reference tetrahedron beak was then transformed to match the orientation of the initial reconstruction. Reconstruction was performed for the initial position before the co- song gesture occurred and the end position after the co-song gesture was finished. To analyze the movement from the initial and end position of the co-song gesture, the two reconstructions were used as inputs into an implementation of the Kabsch algorithm, which calculated principal angle (PA) measurements for each gesture. PA describes the total magnitude of rotation required to align the initial head pose with the end head pose without specifying direction. To account for multiple degrees of freedom (DoF), the contribution values indicate how much each rotation (yaw, pitch, and roll) contributes to the overall movement. Values closer to 1 indicate more pure rotations while values closer to 0 indicate little to no contribution. To examine the effect of social context and syllable complexity on gesture PA, we conducted a linear mixed model analysis (SPSS, 29.0.2.0). To ensure normality of residuals and homogeneity of variance, PA was transformed using a square-root transformation. The model used social context (directed vs undirected song) and syllable complexity score (95% CI method) as fixed effects and included random intercepts for each bird and each syllable using a VC covariance type (*n* = 3 birds, 19 syllables, 660 total gestures/PA scores).

### Social context experiment

The recording and analysis of adult males singing alone (or undirected song) was described previously. To record the co-song gestures under female context (directed song), the housing environment and cage/recording setup was the same as previously mentioned for the song and gesture recording section. In each experiment, each adult male (*n* = 11 birds) was introduced to an adult female (*n* = 10 virgin females) for 10-minute periods. Following the 10 minutes, the female was then removed from the cage, followed by a one-hour isolation period. This process was repeated three times, averaging 30 minutes of interaction between each focal male and female. Video and audio recordings were captured as previously described; approximately 50 motifs and corresponding co-song gestures were coded.

### Deafening study (deprivation of auditory feedback)

Because bilateral deafening is a relatively invasive surgical procedure, we sought to deprive birds of auditory feedback through a less invasive method before resorting to deafening. We first attempted a “white noise experiment,” in which birds were exposed to loud white noise for several months, with the expectation that this would temporarily block auditory feedback and allow us to test the alignment between birdsong and co-song gestures in the absence of auditory feedback. However, this approach proved unworkable, as it was extremely difficult to precisely quantify and align song and gesture under continuous loud white noise. We therefore proceeded with the deafening procedure only after this alternative failed.

For deafening procedure, males were deafened by cochlear extirpation following the protocol from Konishi [56] and a previous study [57]. Briefly, birds were fully anesthetized throughout the entire procedure. After anesthetization with isoflurane, a small incision was made through the back of the skull overlying the semicircular canals. A hooked wire was lowered into this hole and used to pull out the cochlea (n = 5 adult males, age 1–3 years). Immediately after the surgery, animals were placed on a heating pad for recovery, and animals were given buprenorphine (analgesic), 0.05 mg/kg, to relieve any possible pain. Animals were closely monitored daily until restoration of normal behavior and were then returned to their original cages with food and water. No signs of pain, distress, or sickness were observed in post-surgical animals over the course of the experiments (signs of sickness include rapid or slow breathing, weight loss, and puffed feathers).

Two months after recovery, post-surgical birds were placed in a recording chamber to record their song and co-song gestures. Following the conclusion of the experiments, all birds were euthanized and brain tissue was examined to confirm that deafening procedure had been performed correctly.

### Electrolytic lesioning study

Adult males (n=5 birds) who were at least 1 year old were used for this study. For bilateral electrolytic lesions of HVC, insect pins (size 000, Fine Science Tools, CA) insulated with Insl-X were used as electrodes [58]. The electrode end was lowered down, and lesions were made in the rostro-caudal range 1 mm at 50 uA each for 40 seconds. For sham control birds (n=5 birds), the wire was lowered down but no electrical current was delivered. Post-surgical birds were fully anesthetized throughout the entire procedure. Immediately after the surgery and during the post-surgery period, animals were given buprenorphine, 0.05 mg/kg, to relieve any possible pain.

Animals were placed on the heating pad for recovery, and they were closely monitored daily until restoration of normal behavior and then were placed to original cages for audio-video recordings.

After the conclusions of the experiments, all the post-lesioned birds were euthanized, and we examined the brains of these post-surgical birds to confirm that the lesioned sites were precisely targeted.

### Playback experiment

Each adult virgin female (*n* = 8 females, approximately 1 year old) was kept in a soundproof chamber with visual and acoustic isolation from other birds and was allowed to acclimate to the cage for 3 days before experiments began. Each female was shown an adult male singing (n= 7 male birds) on two screens at either end of the cage. During the playback experiment, videos were played via an iPad screen on both the left and right sides of the cage (cage size: 25.5 × 30.5 × 33 cm; **Supplementary Figure. 5**). Birds were supplied water, seeds, eggs, and gravel.

The type of video, normal or delayed, alternated sides between experimental trials to control for side preferences. Each screen had a perch directly in front of it, and time spent on the perch while watching the screen was quantified. The iPad screens were kept at equal brightness and a sound level of 85 dB. Screens were adjusted to be the same height and were level with the perches. Finch behavior was video recorded (Reolink E1 Zoom) and later quantified (BORIS).

Two independent coders watched video recordings of the trials and selected the video frame where the finch started and stopped watching the screen. The total time was then quantified from the time fragments obtained. Experimental females received a 10-minute break between each video exposure trial, and finches watched all 4 strangers on each test day. Finches followed this procedure for 2 days.

## Acknowledgments

We thank the animal caretakers at Colgate University for providing excellent animal care. We are grateful for all of the Colgate students who assisted in this project, including Alexis Romero, Bella Jaffe, Chidinma Okafor, Juhyun Park, Isla Gao, Cailen Geller, Grace Ciaravino, Allan Crounse, Kajol Luplunge, Ana Diaz, Caitlin Mooney, and Zhuangyi Liao. This research is supported by Colgate University’s Picker ISI major research grant, MBBI major grant, and Research Council Picker research grant.

## Supplementary Video#1

An example of co-song gestures in the zebra finch. These gesture movements are primarily lateral head rotations dominated by yaw, and they are rhythmic movements that accompany each song motif. The video was created by a trained DeepLabCut mode to generate 2D coordinates for key points of the beak (colored dots to represent crown, left base corner, right base corner, and tip).

---

**Supplementary Figure 1.**
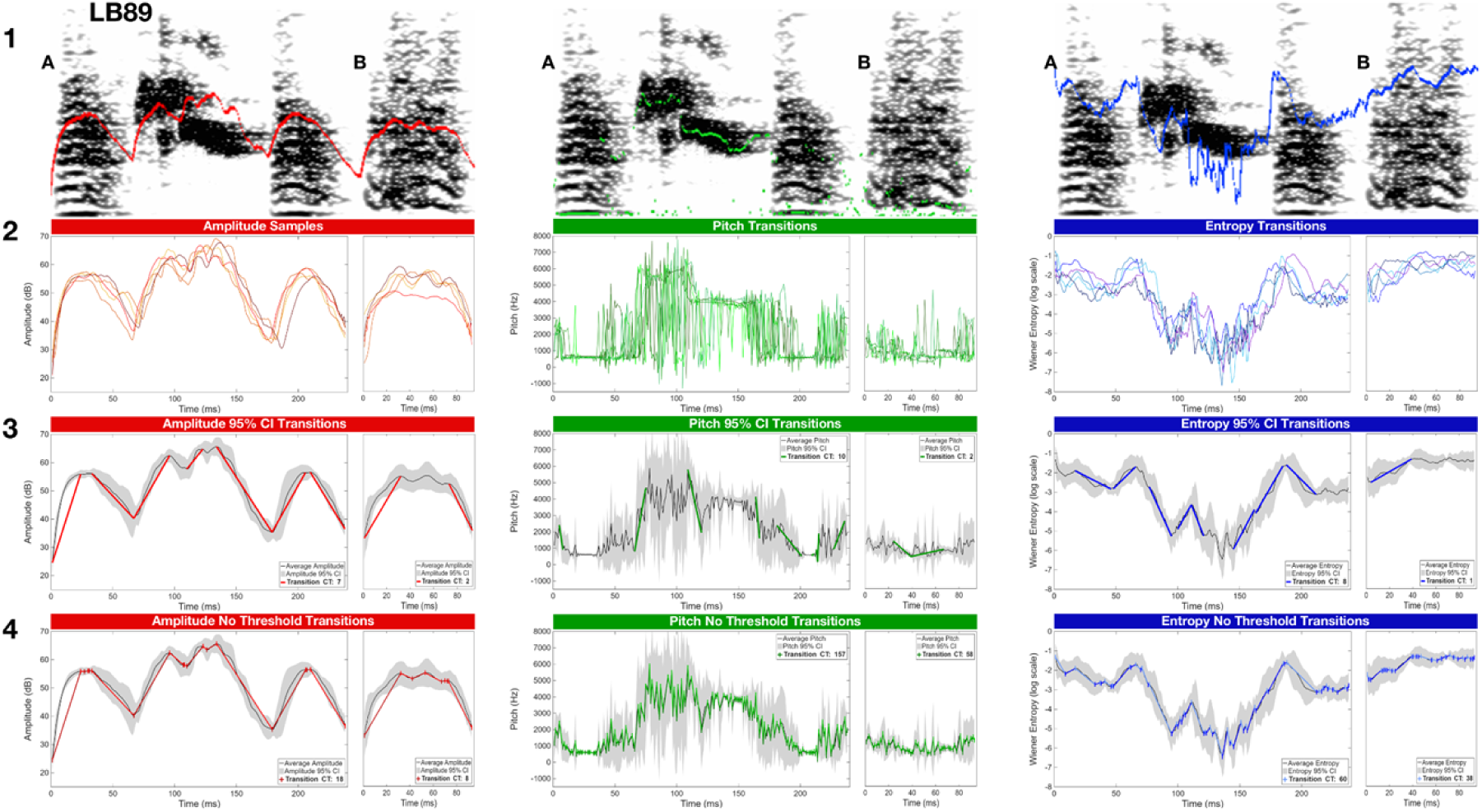
Both accumulative significant transitions and total transitions in acoustic features quantify syllable complexity. Example of acoustic transition quantification process. **(1)** Acoustic feature measurements at each timepoint within a syllable interval in SAP (amplitude, pitch, and entropy for syllables A and B of LB89 over time illustrated). **(2)** Five repetitions of each syllable were exported to MATLAB and resampled to match the averaged duration for that syllable. Local maxima and minima for each acoustic feature’s average line were calculated and intervals between them considered as potential transitions. **(3)** Result of 95% CI approach to transition quantification; shortest time intervals with start and end points that did not have overlapping CIs were selected as significant transitions in an acoustic feature. **(4)** Result of the no-threshold approach to transition quantification; all time intervals between adjacent local maxima/minima were counted as transitions in an acoustic feature.

**Supplementary Figure 2.**
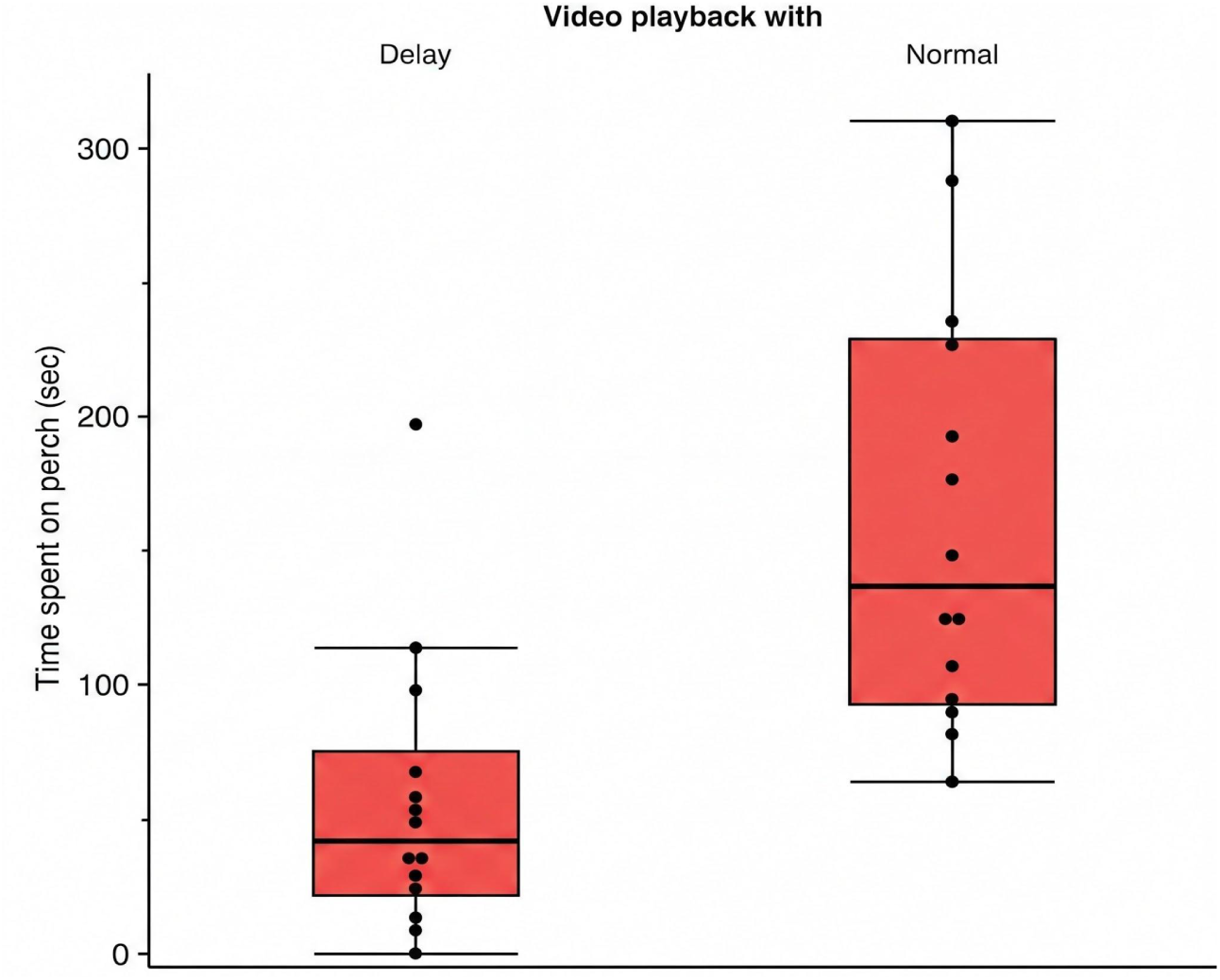
Females prefer rhythmically matched songs and gestures. We found that females (*n* = 7 birds, each bird had 2 trials) spent significantly more time on the perch near normal, synchronous playback, suggesting the females can identify the mismatching of co-song gestures. Each dot represents a trial of a female.

**Supplementary Figure 3.**
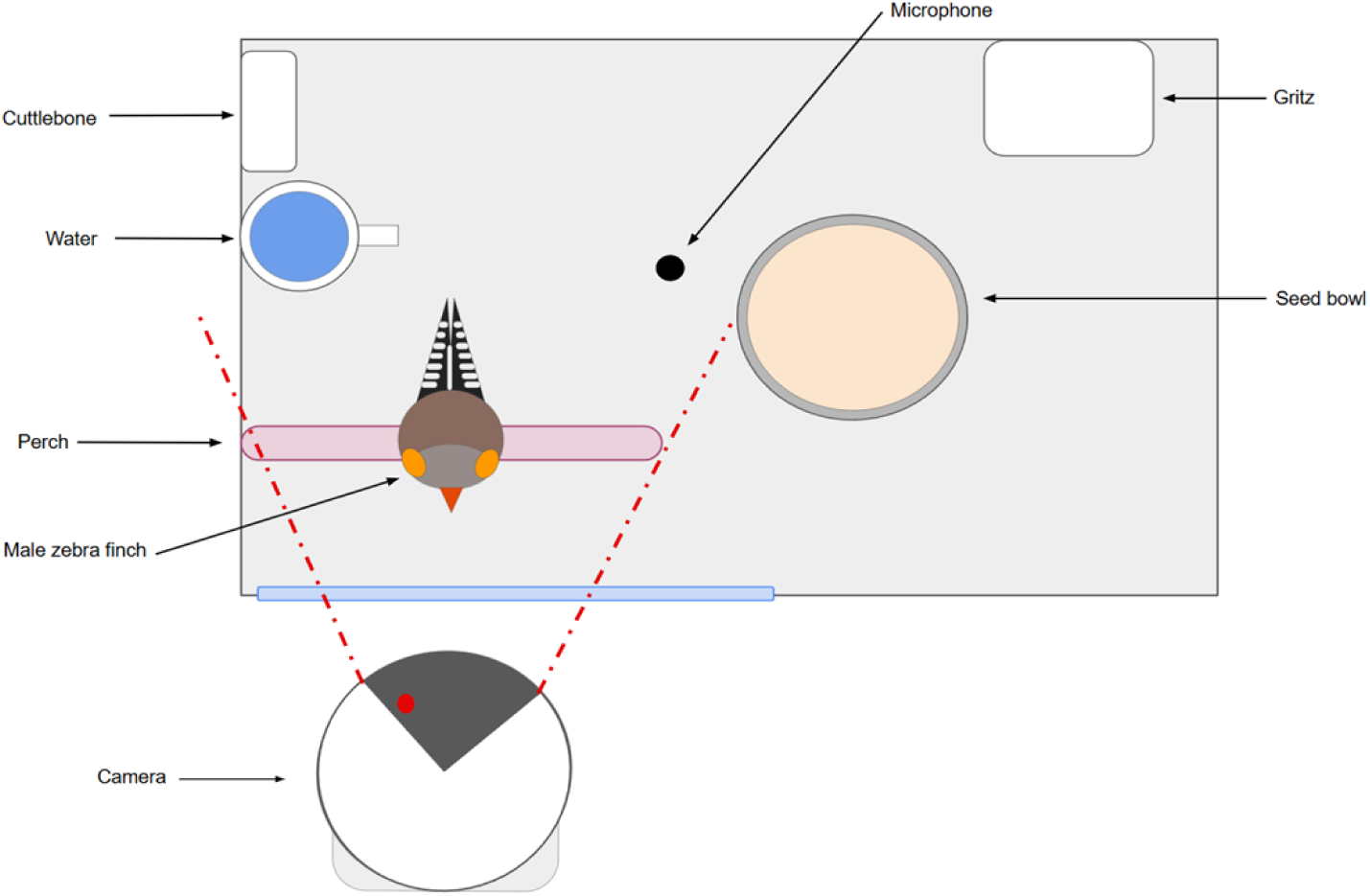
Experimental recording setup for behavioral coding and song analysis. Example cage setup for recording of undirected song of adult male zebra finches. Seeds, chicken egg supplements, water, grits, cuttlebone, perches, and enrichment toys (not pictured, varied by cage) were provided. A WiFi-connected Reolink E1 Zoom or Reolink 4MP camera obtained video footage; a condenser microphone (Audio Technica, AT801) obtained audio recordings.

**Supplementary Figure 4.**
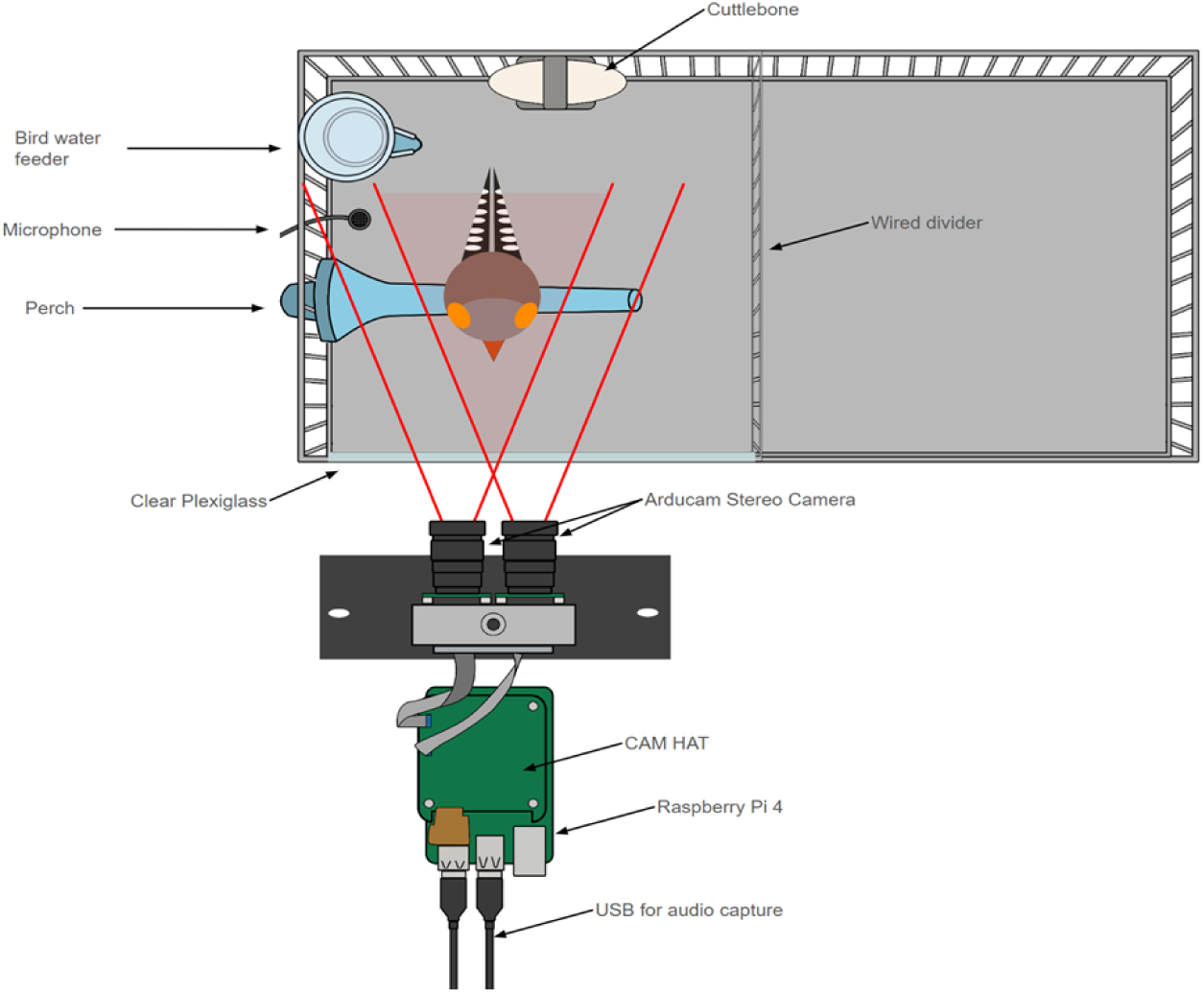
Experimental recording setup for 3D reconstruction model. Example cage setup for recording of male zebra finches to create a 3D reconstruction of directed and undirected co-song gestures. Seeds, chicken egg supplements, water, grits, cuttlebone, perches, and enrichment toys (not pictured, varied by cage) were provided. 2 12.3MP 477P Raspberry Pi Arducam cameras and a Behringer U Phoria UMC202HD USB Audio Interface (not pictured) with a condenser microphone (Audio Technica, AT801) were connected to a Raspberry Pi 4 to capture audio and video. A wired divider was used to ensure males were visible in the camera.

**Supplementary Figure 5.**
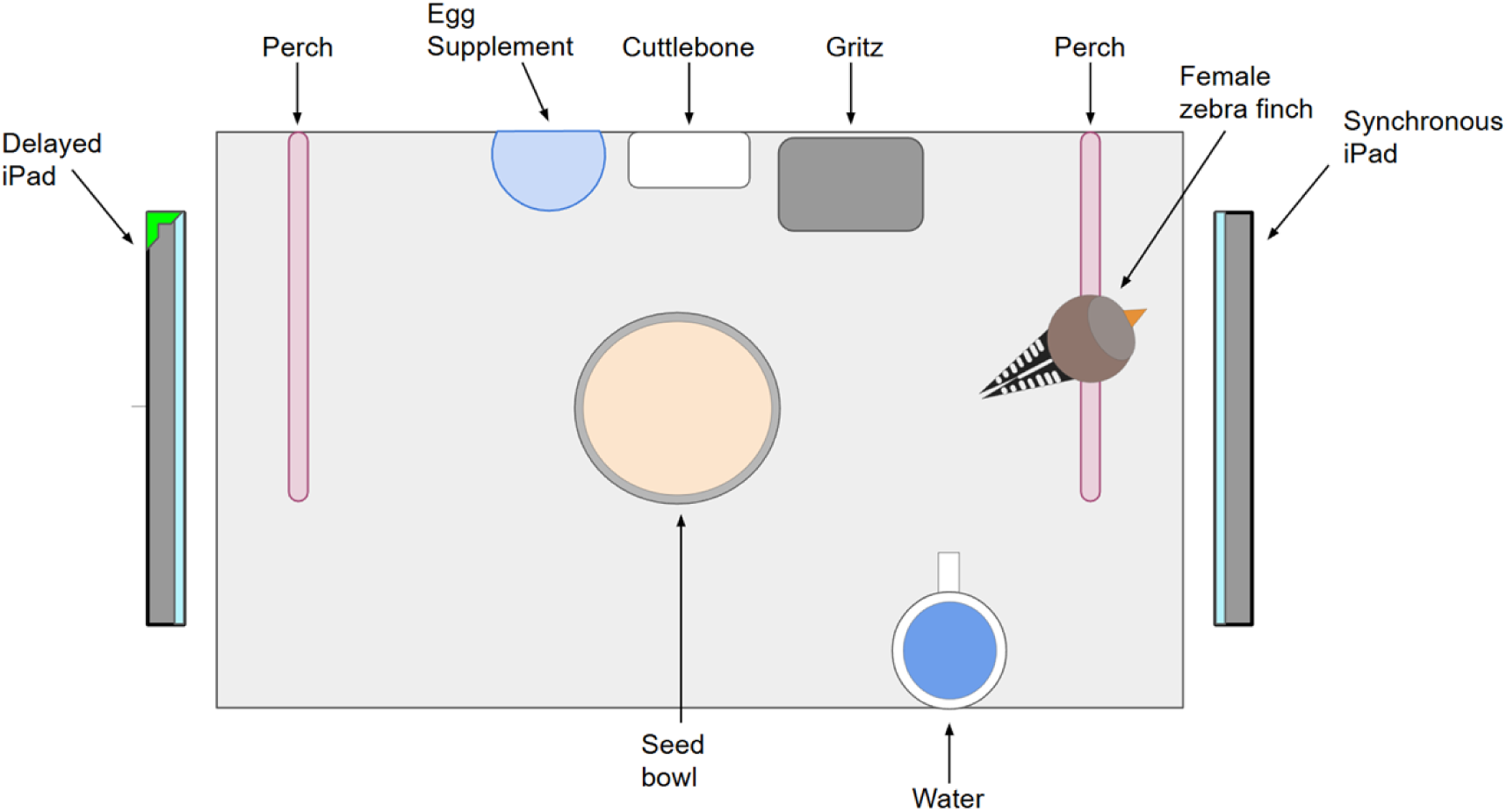
Experimental recording setup for playback procedure. Example cage setup for recording of female zebra finch behavior towards playback of male undirected song. Seeds, chicken egg supplements, water, grits, cuttlebone, two perches, and enrichment toys (not pictured, varied by cage) were provided. Two iPads were present at each end of the cage, one playing a recording of male undirected song with synchronous video and audio, while the other playing asynchronous audio and video (delayed audio; mismatched song and co-song gestures). Which iPad had synchronized audio and video was randomized for each trial. A WiFi-connected Reolink E1 Zoom camera obtained aerial video footage of the bird’s position in the cage. Time spent on each perch was recorded.

